## Supplementary Figures 1-15 for "Noncaloric monosaccharides induce excessive sprouting angiogenesis in zebrafish via foxo1a-marcksl1a signal"

**Running title: Noncaloric Monosaccharides Induce Excessive Angiogenesis**

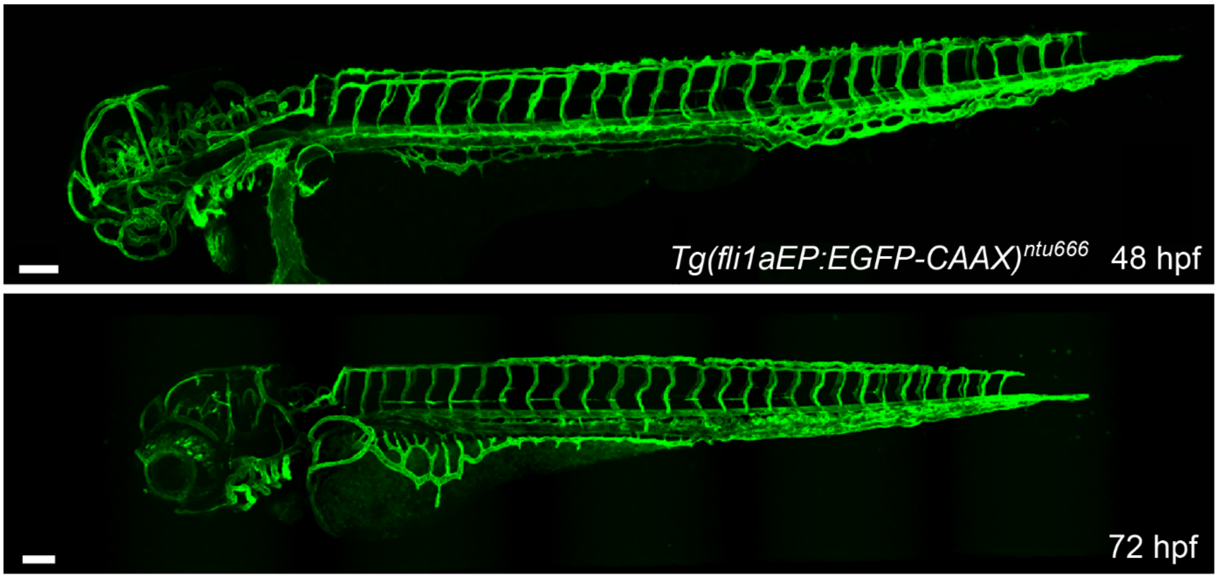


**Supplementary Figure 1 Confocal imaging analysis of *Tg(fli1aEP:EGFP-CAAX)*$^{ntu666}$ embryos at 48 hpf (a) and 3 dpf (b). Scale Bar=100 µm.**

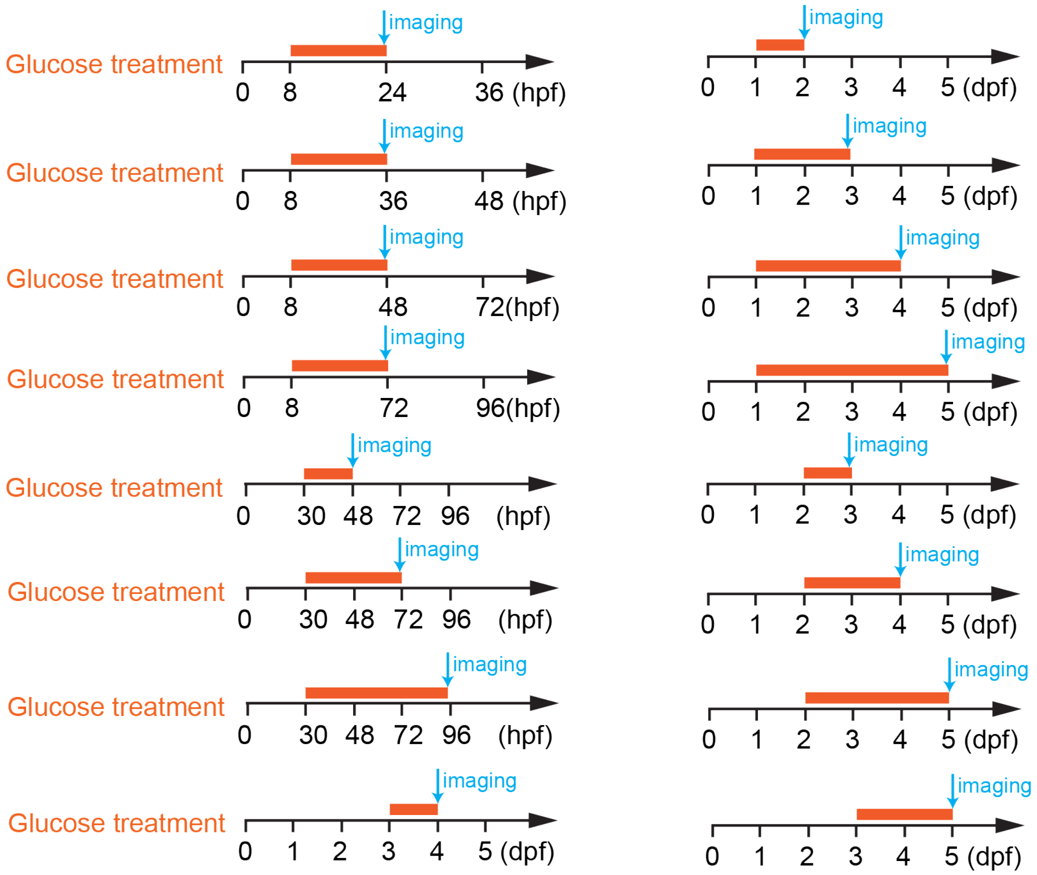


**Supplementary Figure 2 The diagrams show the glucose treatment time window and imaging time point.**

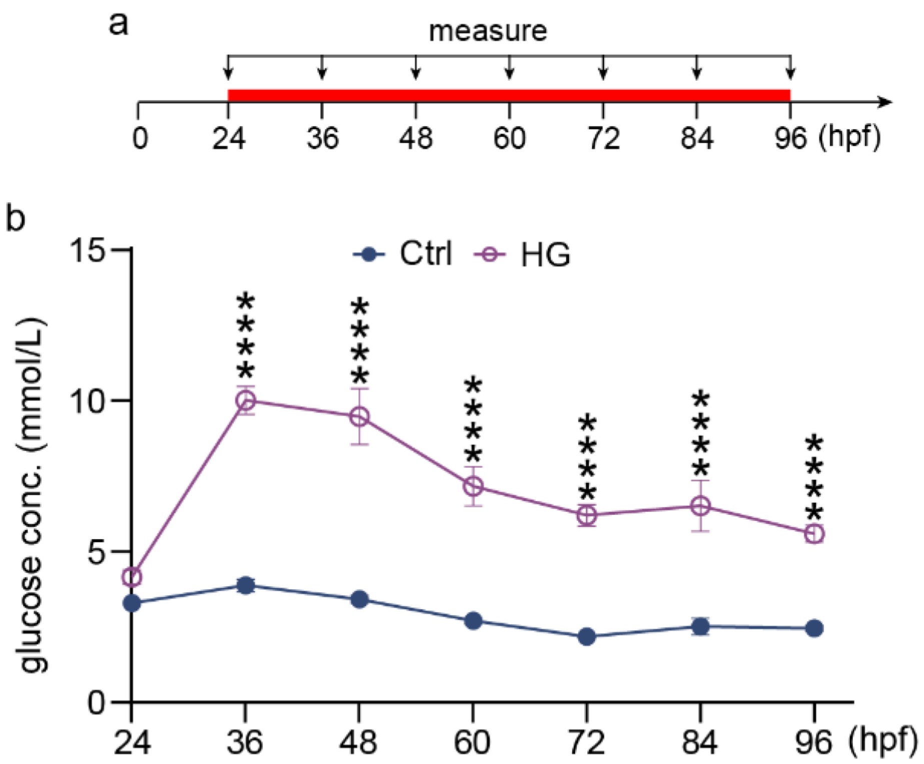


**Supplementary Figure 3 Total glucose concentrations at different development stages in control and high glucose treated embryos. a**, A diagram showing the glucose treatment time window and concentration measuring time point. **b**, Statistical analysis of the glucose concentration in control and high glucose-treated embryos. one-way ANOVA, ****p<0.0001.

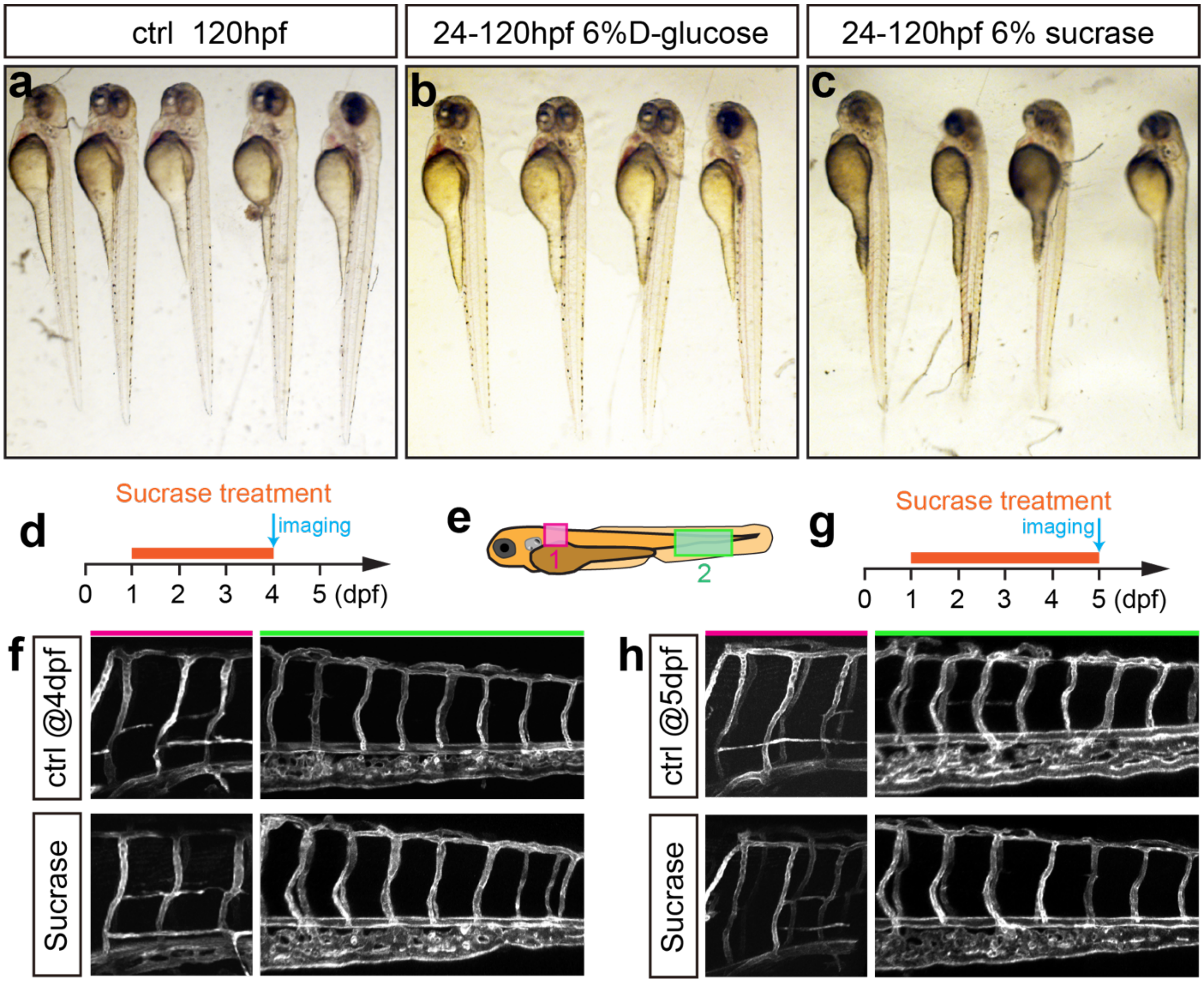


**Supplementary Figure 4 Stereo microscopic analysis of control, glucose, and sucrase-treated embryos in a bright field. a-c**, Imaging analysis of control, glucose and sucrase treated embryos in bright field. **d**, A diagram showing the sucrase treatment time window and imaging time point. **e**, A diagram indicating the imaging positions of the zebrafish embryos. **f,** Confocal imaging analysis of the control and sucrase treated *Tg(fli1aEP:EGFP-CAAX)*$^{ntu666}$ embryos. The red bar indicates position 1; the green bar indicates the position. **g**, A diagram showing the sucrase treatment time window and imaging time point. **h**, Confocal imaging analysis of the control and sucrase treated *Tg(fli1aEP:EGFP-CAAX)*$^{ntu666}$ embryos. The red bar indicates position 1; the green bar indicates the position.

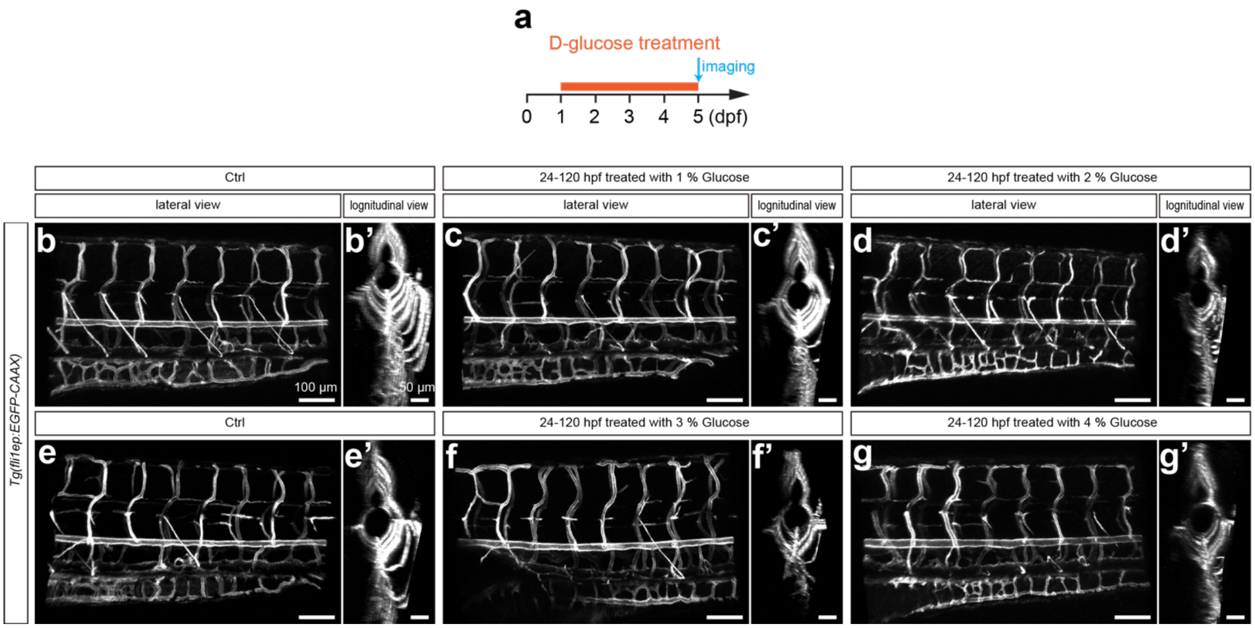


**Supplementary Figure 5 Confocal imaging analysis of 1%4% glucose-treated blood vessels. a, A diagram showing the glucose treatment time window and imaging time point. b—g', Confocal imaging analysis of the control, 1%, 2%, 3%, and 4% glucose-**treated *Tg(fli1aEP:EGFP-CAAX)*$^{ntu666}$ embryos.

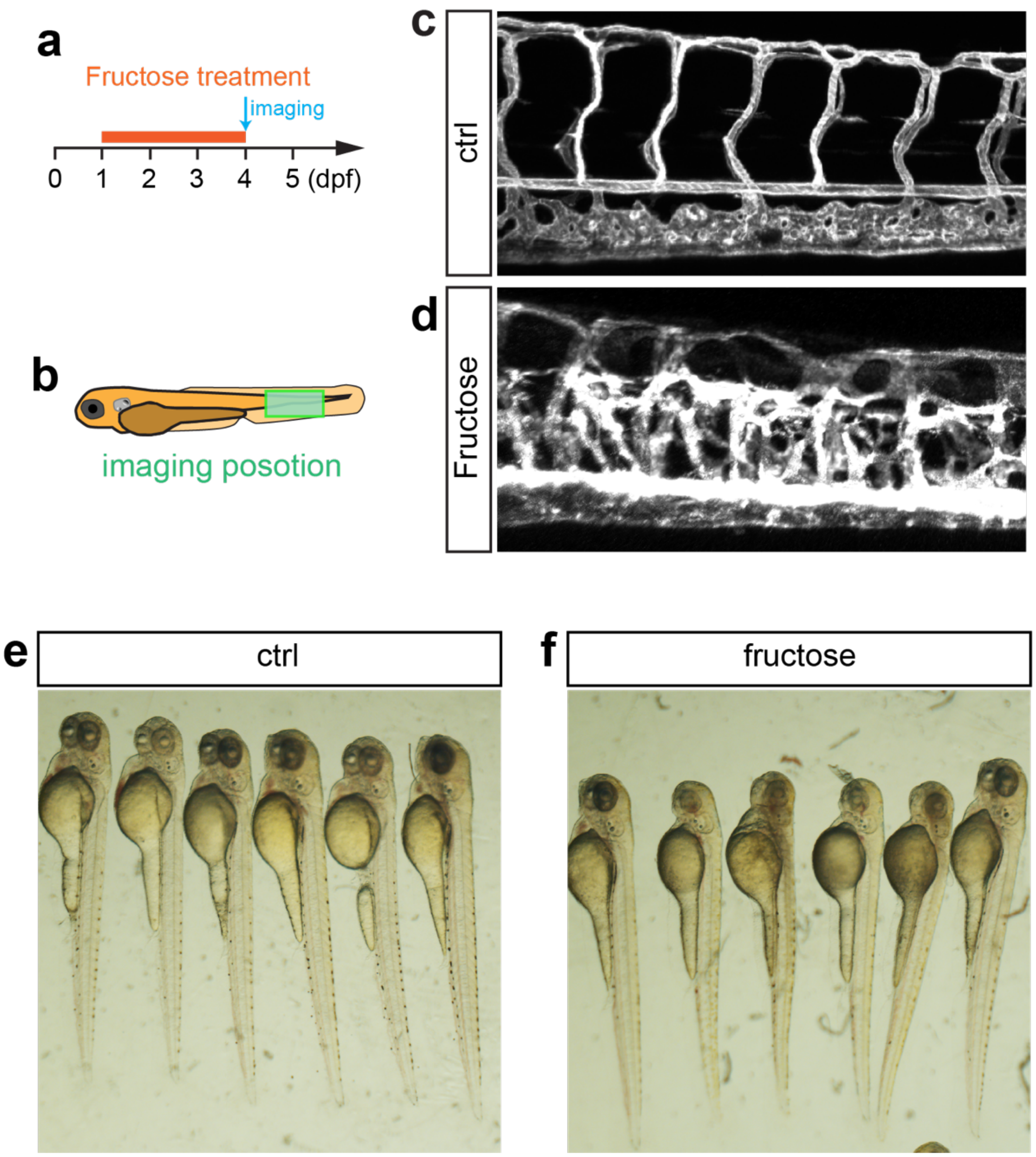


**Supplementary Figure 6 Fructose treatment caused excessive angiogenesis in zebrafish. a**, A diagram showing the fructose treatment time window and imaging time point. **b**, A diagram indicating the imaging position of the zebrafish embryos. **c-d**, Confocal imaging analysis of the control and glucose-treated Tg(fli1aEP:EGFP-CAAX)ntu666 embryos. e-f, Imaging analysis of control and fructose-treated embryos in bright field.

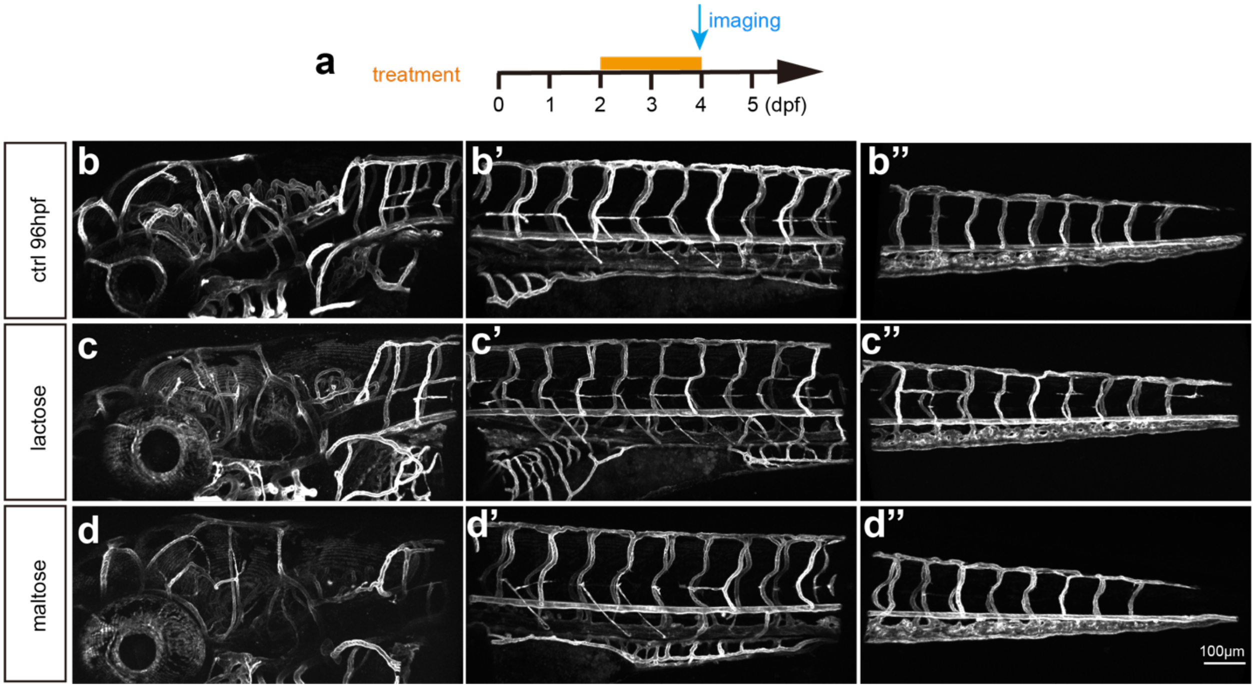


**Supplementary Figure 7 Lactose and maltose treatment did not cause excessive angiogenesis in zebrafish. a**, A diagram showing the lactose and maltose treatment time window and imaging time point. **b-d''**, Confocal imaging analysis of the control, lactose, and maltose-treated *Tg(fli1aEP:EGFP-CAAX)*$^{ntu666}$ embryos.

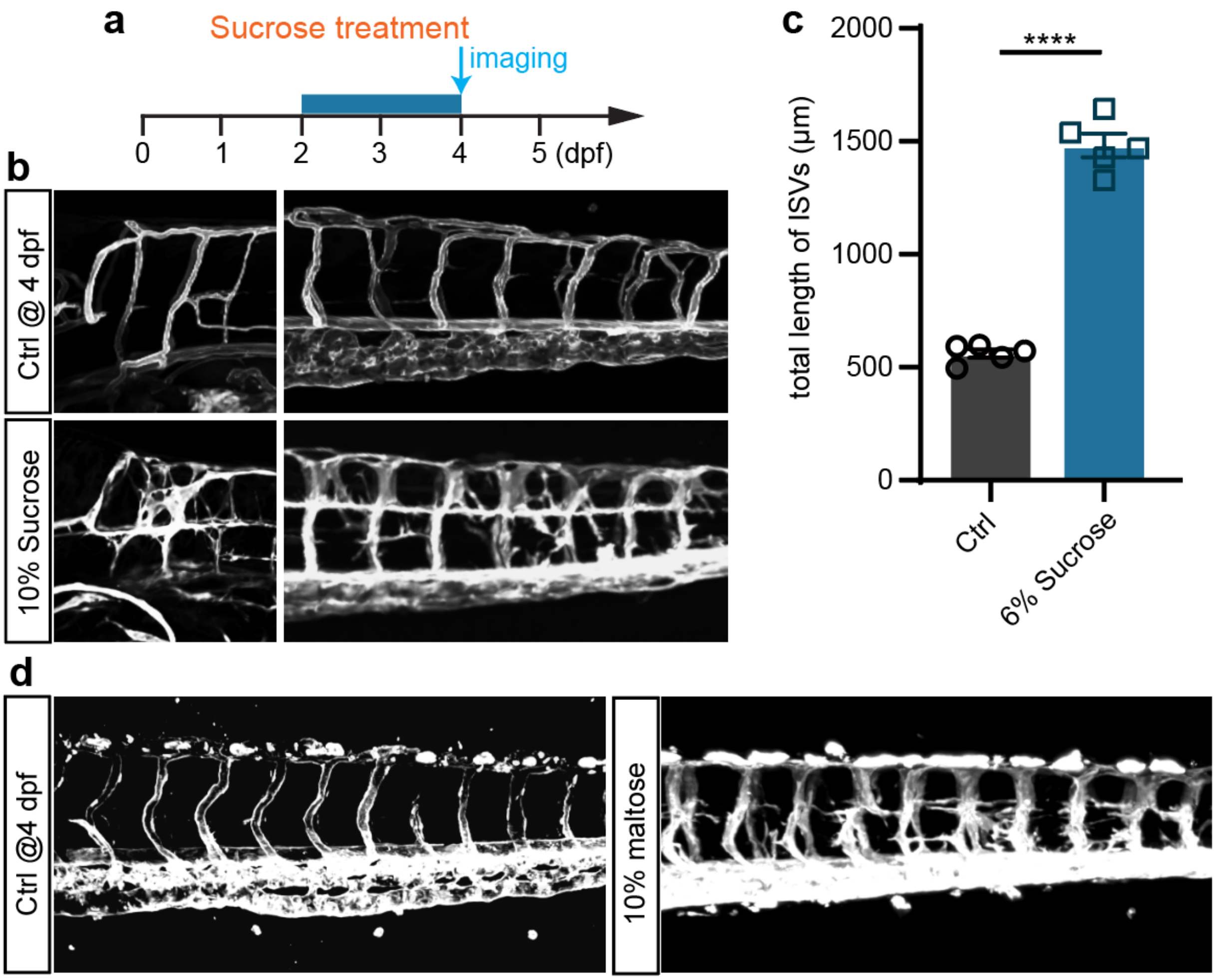


**Supplementary Figure 8 Higher concentration sucrase and maltose treatment cause excessive angiogenesis in zebrafish embryos. a**, A diagram showing the sucrase and maltose treatment time window and imaging time point. **b,** Confocal imaging analysis of the control and sucrase treated *Tg(fli1aEP:EGFP-CAAX)*$^{ntu666}$ embryos. **c,** Statistical analysis of the total length of ISVs in control and sucrase-treated embryos. *t-test*, *****p*<0.0001.**d,** Confocal imaging analysis of the control and maltose-treated *Tg(fli1aEP:EGFP-CAAX)*$^{ntu666}$ embryos.

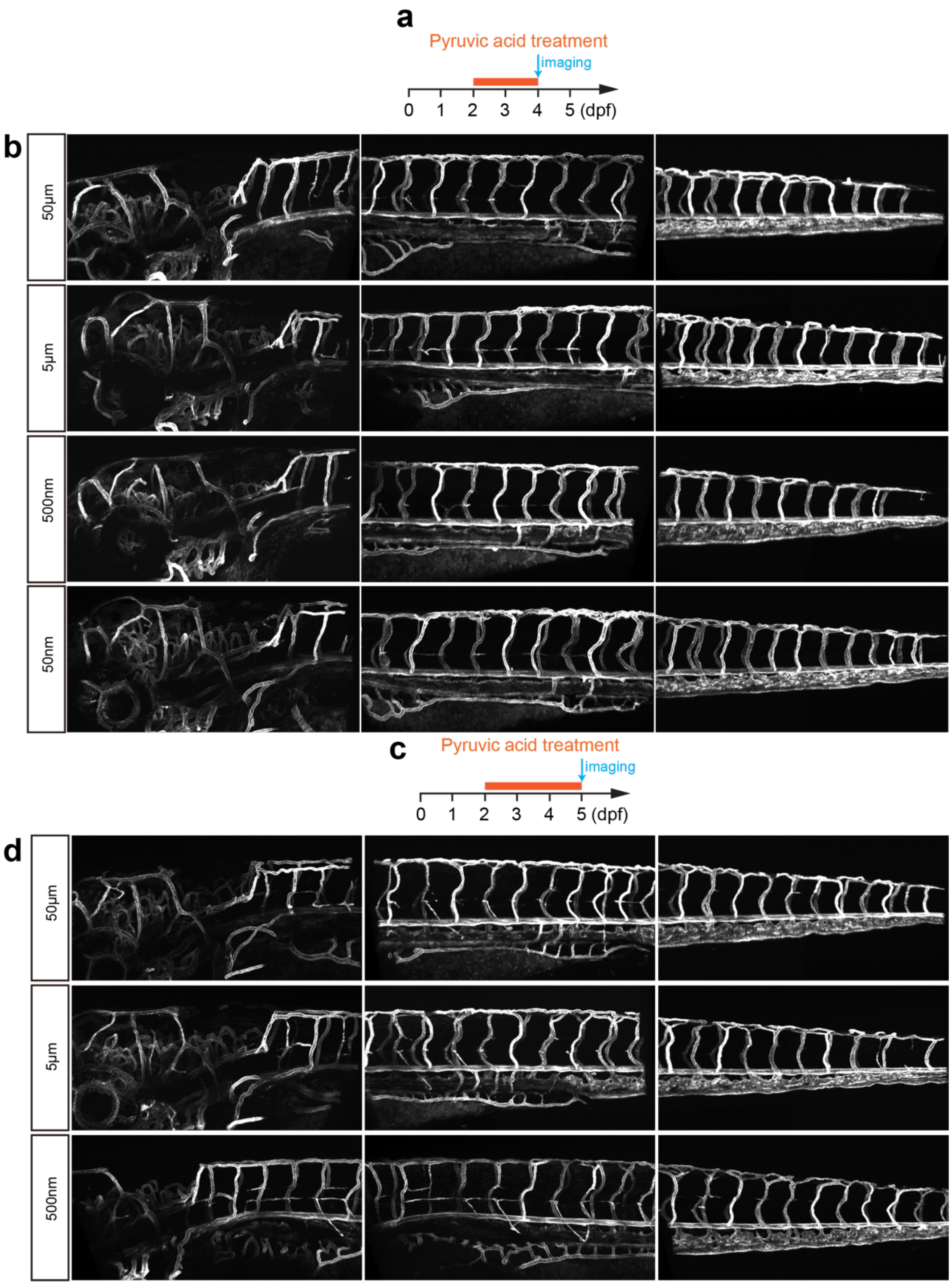


**Supplementary Figure 9 Pyruvic acid treatment did not cause excessive angiogenesis in zebrafish. a**, A diagram showing the pyruvic acid treatment time window and imaging time point. **b**, Confocal imaging analysis of the pyruvic acid treated *Tg(fli1aEP:EGFP-CAAX)*$^{ntu666}$ embryos. **c**, A diagram showing the pyruvic acid treatment time window and imaging time point. **d**, Confocal imaging analysis of the pyruvic acid treated *Tg(fli1aEP:EGFP-CAAX)*$^{ntu666}$ embryos.

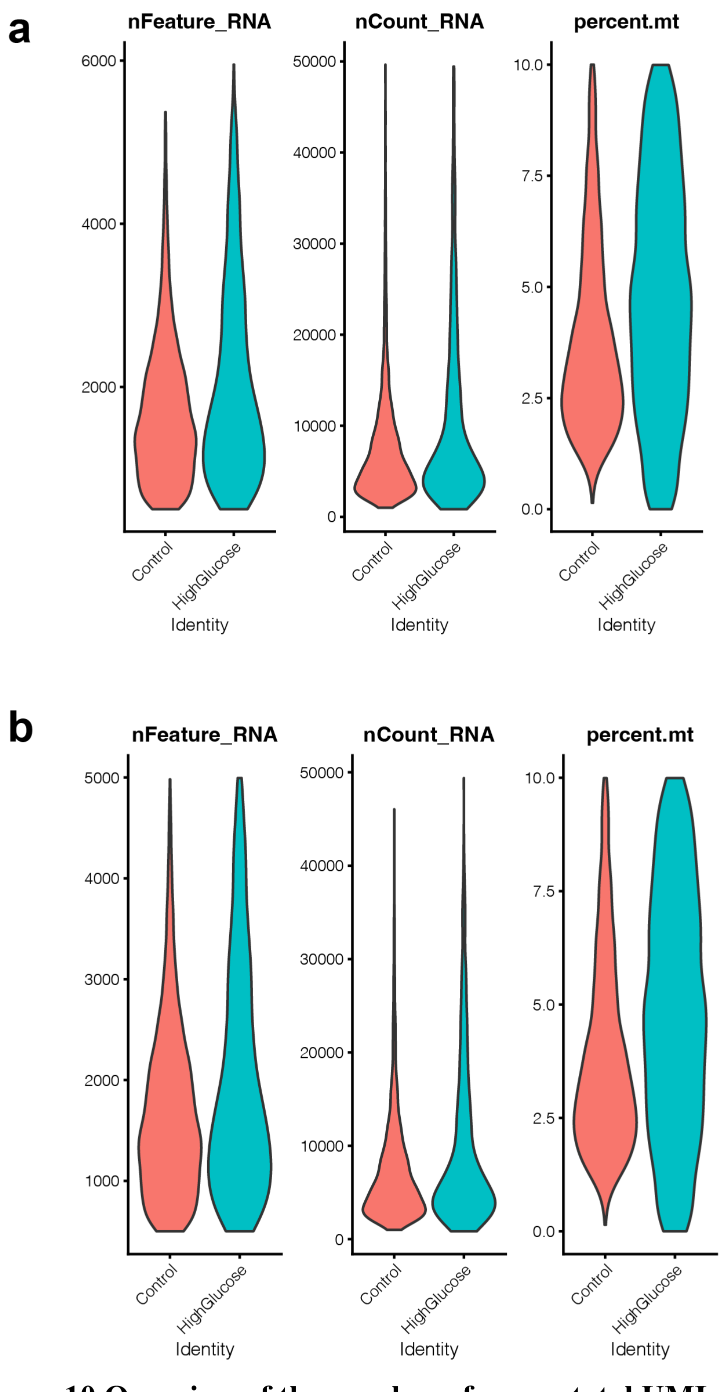


**Supplementary Figure 10 Overview of the number of genes, total UMIs and percentage of mitochondrial UMIs for the single cell RNA sequencing. a**, Before filtering. **b**, After filtering. Cell selection criteria: 500< number of genes < 3000; 0< percentage of mitochondrial UMIs < 5%.

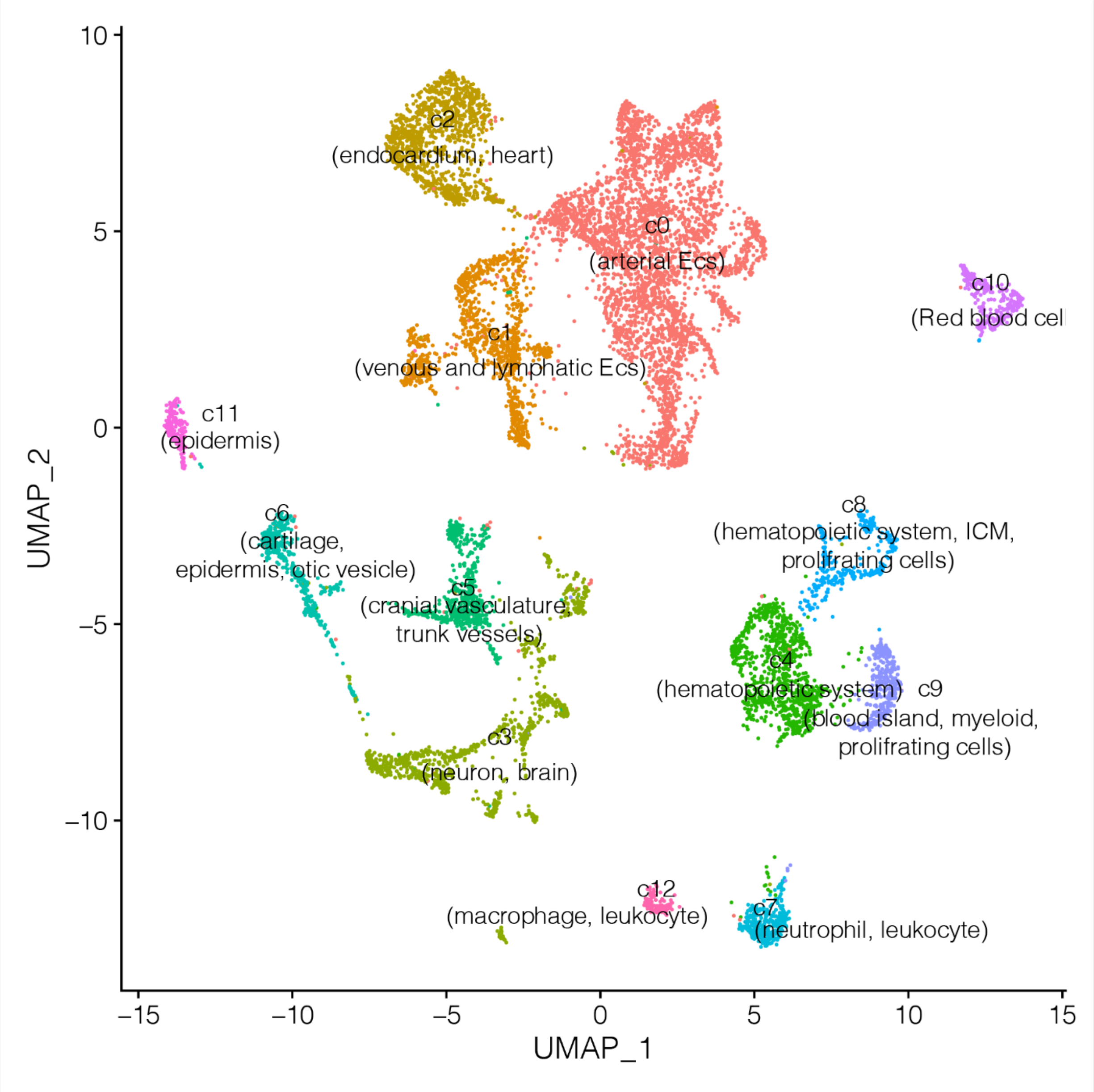


**Supplementary Figure 11 UMAP representation of EC subpopulations. All single cells (after filtering) from control and l- glucose treated were included in this illustration.**

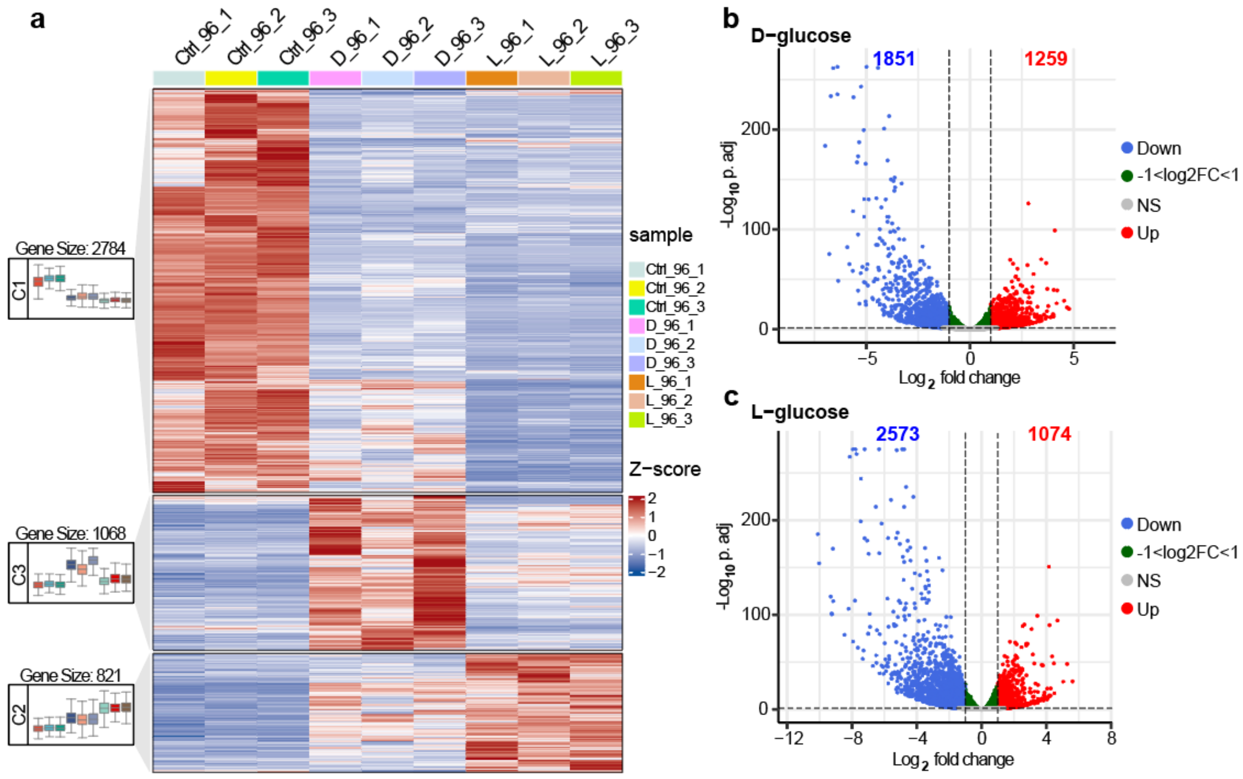


**Supplementary Figure 12 Transcriptome sequencing analysis of control, high D-glucose, and high L-glucose treated embryos. a**, The heatmap of differentially expressed genes (DEGs). **b**, Volcano map of DEGs after high D-glucose treatment. **c,** Volcano map of DEGs after high L-glucose treatment.

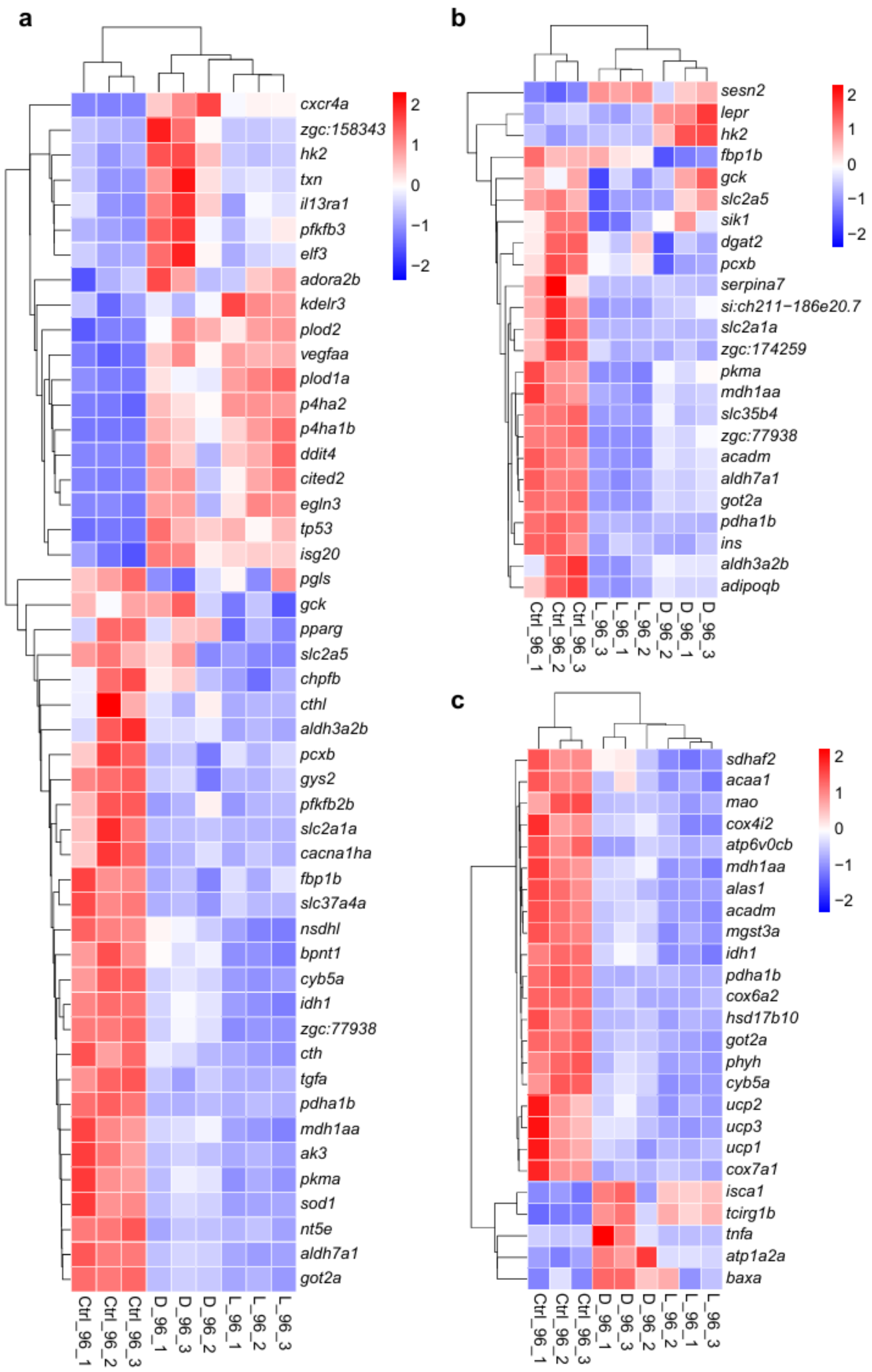


**Supplementary Figure 13 The heatmap of metabolism related genes. a,** The heatmap of gluconeogenesis related genes. **b,** The heatmap of glycolysis related genes. **c,** The heatmap of oxidative phosphorylation related genes.

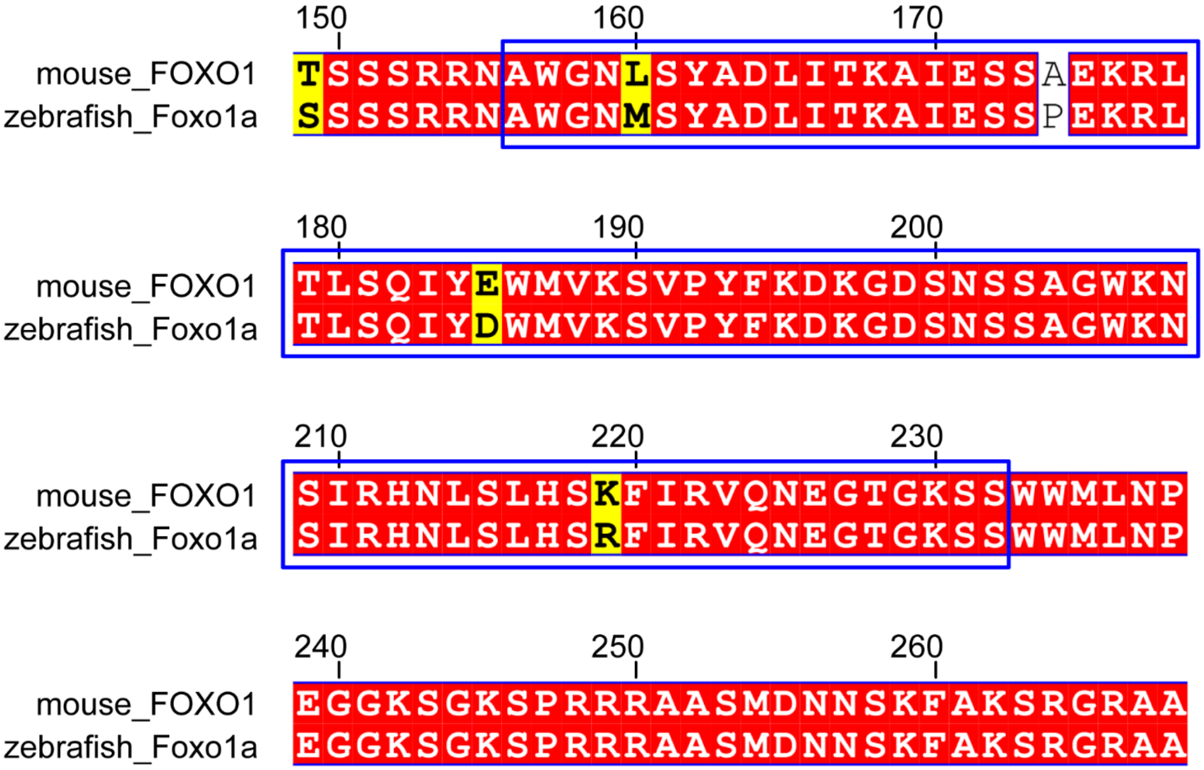


**Supplementary Figure 14 Multiple amino acid sequence alignment of mouse FOXO1 and zebrafish Foxo1a.** The DNA binding domain of mouse FOXO1 is demarcated with blue.

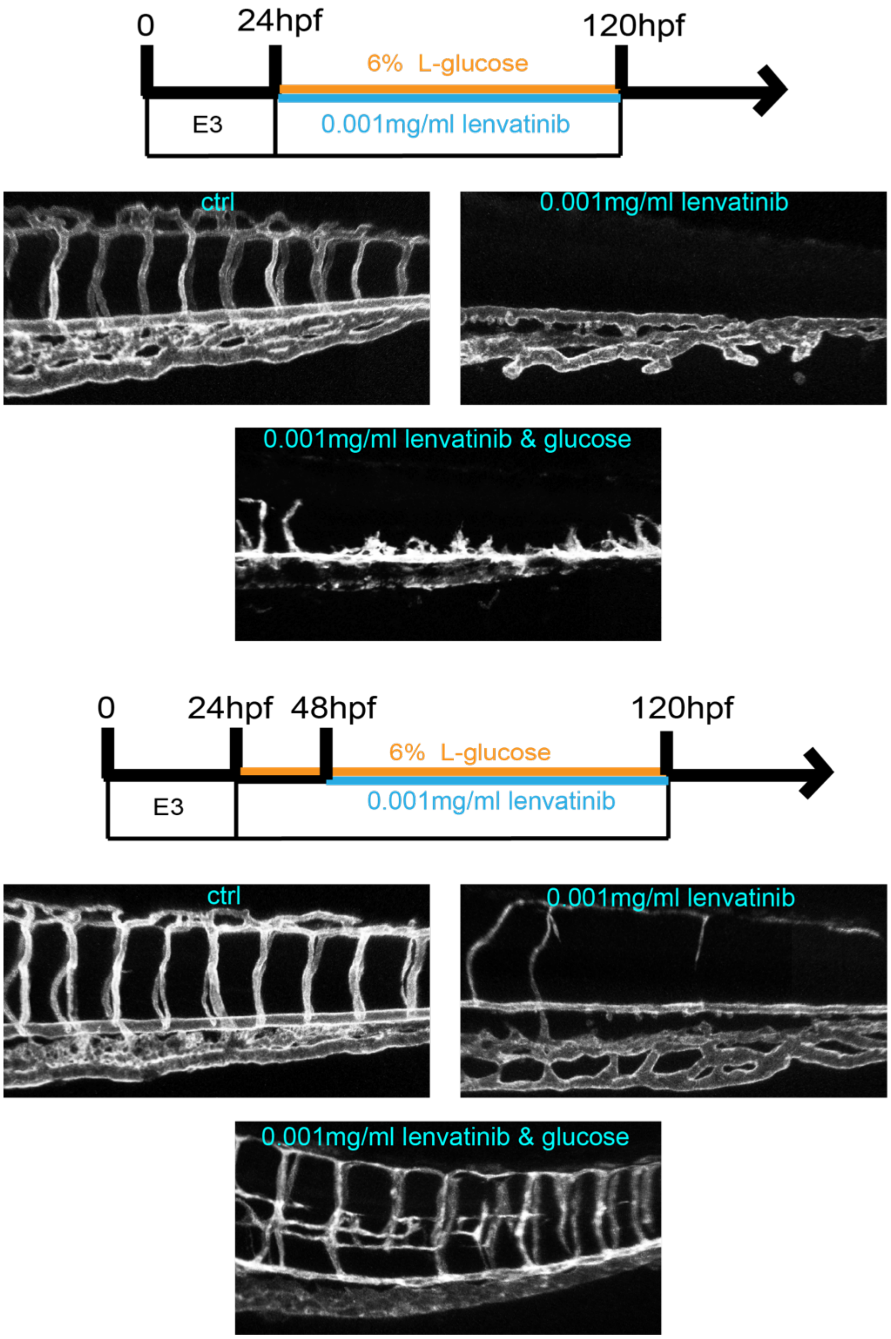


**Supplementary Figure 15 Confocal imaging analysis of blood vessels in the embryos with lenvatinib treatment. a**, A diagram showing the lenvatinib treatment time window and imaging time point. **b**, Confocal imaging analysis of the control and lenvatinib treated *Tg(fli1aEP:EGFP-CAAX)*$^{ntu666}$ embryos. **c**, A diagram showing the lenvatinib treatment time window and imaging time point. **d**, Confocal imaging analysis of the control and lenvatinib treated *Tg(fli1aEP:EGFP-CAAX)*$^{ntu666}$ embryos.
